# High Glucose Diet Induces Hepatic Iron Overload Contributing to Metabolic Dysfunction

**DOI:** 10.1101/2025.03.20.644441

**Authors:** Amanda Caceres, Nathaniel H.O. Harder, Jacob P. Padilla, Samuel E. Janisse, Austin M. Cole, Sonia E. Roedersheimer, Marie C. Heffern

## Abstract

Iron is an essential biometal, critical in processes that include oxygen transport, mitochondrial respiration, and cell signaling. Iron dyshomeostasis is linked with hyperglycemia and associated metabolic disorders, but the underlying mechanisms are poorly understood. To investigate these mechanisms, we conducted a short-term, four-week, in vivo study on mice given water supplemented with glucose. The short time frame was sufficient to cause metabolic shifts in the liver towards triglyceride synthesis. We sought to comprehensively track iron trafficking by analyzing liver and serum markers of iron metabolism alongside LC-ICP-MS analysis of iron speciation, which is a new approach in this context. Glucose supplementation induced changes in iron regulation despite equal dietary iron intake between groups. Specifically, we observed increased uptake of transferrin-bound iron from the serum and an iron overload state in the liver. We developed and applied a cell-based models of this glucose-induced iron overload state and found that, on the one hand, the anti-diabetic drug metformin could restore iron regulation; on the other hand, the iron chelator, deferoxamine, could restore glucose metabolism. Taken together, our studies reveal that early hyperglycemia is sufficient to cause disruptions in iron regulations, pointing to iron overload as viable therapeutic target in metabolic dysfunction.

## INTRODUCTION

Hyperglycemia is known to occur in a variety of disease states including metabolic diseases like diabetes and metabolically associated fatty liver disease, cancer, stroke, and acute illness. In all cases, the occurrence of hyperglycemia leads to worse disease outcomes^1–5^. Obesity and diabetes incidence rates are increasing in the United States^6^. Dietary changes, in particular the Western diet which is high in sugars and fats, are thought to be major contributors to the increased prevalence of obesity and diabetes^7^. Obesity is the biggest predictor for hyperglycemic state, however, studies tend to focus on insulin resistant diabetic disease state while early disease states remain understudied. Additionally, many studies focus on a combination of long-term high-fat and high-sugar diets, however, the role of dietary sugars alone in disease pathology is not understood^8–12^.

Metal micronutrients are an essential dietary component and have been linked to several disease states. In particular, iron homeostasis has been tied to both metabolically associated fatty liver disease and diabetes^13–17^. It is thought that iron mediates disease onset by affecting glucose metabolism, insulin secretion, and insulin sensitivity^13–15,18,19^. Iron is an essential cofactor for fuel oxidation, yet its redox activity and potential to cause oxidative damage requires that it remain tightly regulated. The links between iron and metabolism are well documented. Iron uptake has been found to increase when needed for fuel oxidation while glucose and ethanol metabolism are dependent on the availability of iron^14^. However, many questions remain regarding how dysregulated iron and iron overload interface with metabolic dysfunction.

Previous work by our group had shown effects of short-term consumption of high sugar drinks on clinical parameters of metal micronutrient status in healthy human subjects^20^. Results from this study suggested that even with short-term intake, glucose consumption can impact transition metal metabolism, including effects on iron regulatory proteins. Building on this, our current study aims to better understand the relationship between glucose consumption and iron regulation. We provided male C57/BL6J mice with glucose-supplemented water over a four-week period, and analyzed changes to extracellular serum markers and liver metal stores. By using a combination of immunoassays, quantitative PCR, and inductively coupled plasma mass spectrometry, we were able to both identify and quantify perturbations in systemic iron regulation. We found that a hyperglycemic state increases the uptake of transferrin-bound iron into hepatic tissues, resulting in elevated liver iron stores and decreased serum transferrin saturation. Complementary experiments in cell-based models revealed that iron chelation reduces glucose uptake in hepatic cells which was similarly seen in treatment with the anti-diabetic drug metformin. These results reinforce a link between iron availability and glucose metabolism and further highlight the need to better understand their interplay in disease pathology.

## METHODS

### Chemicals

Ferrous ammonium sulfate, deferoxamine (DFO), metformin hydrochloride, D-glucose, NaCl, SDS, BSA, blotting grade blocker, protease inhibitor, and phosphatase inhibitor were purchased from Millipore Sigma. Tris-HCl, bovine serum albumin fatty acid-free powder (BSA), nitric acid, methanol, nitric acid (TraceMetal Grade), and hydrochloric acid (TraceMetal Grade) were purchased from Fisher Scientific. Blotting grade blocker and β-mercaptoethanol were purchased from Bio-Rad Laboratories. All buffered solutions, metal salt, sugar, and chelator solutions were created using Direct-Q 3 deionized water (>18 MΩ, Millipore).

### Experimental Models

All protocols were approved by the UC Davis Institutional Animal Care and Use Committee and follow the National Research Council’s Guide for the Care and Use of Laboratory Animals. C57BL/6J mice (male, 7-weeks of age) were purchased from Jackson Laboratories (Strain #:000664). Mice were acclimated after shipping for 7 days before beginning experiments. Mice maintained at 20–23 ºC, 45– 65% relative humidity and a light cycle of 14 h light/10 h dark. Mice were fed a standard diet (LabDiet 5001) and maintained with no more than 5 mice per cage.

HepG2 cells were obtained from the UC Berkeley Cell Culture Facility. Cells were grown in complete DMEM media (31053036, Thermo Fisher Scientific) with 10% Avantor Seadigm premium grade fetal bovine serum (97068-085, VWR), 2mM L-glutamine (25-030-081, Gibco), and 1mM sodium pyruvate (11360070, Thermo Fisher Scientific). Cells were maintained in a humidified environment at 37°C and 5% CO_2_ in an incubator and passaged at 70% confluence every 3 to 4 days. All experiments were performed on cells between passages 10 and 20. Sterile culturing and assay plates were used for all experiments.

### High Glucose Diet

Mice were given a high glucose diet through the addition of 10% glucose supplemented into their water. Mice had access to high glucose water and food ad libitum. Mice were weighed weekly and serum was collected by retro-orbital bleed under isoflurane bi-weekly. After four weeks, mice were sacrificed by cardiac puncture under isofluorane and organs were harvested. Organs were flash frozen in liquid nitrogen and stored at -80°C. Mouse serum was collected by centrifugation of cardiac puncture blood after coagulation at 1000 x g for 10 minutes in order to remove red blood cells. After centrifugation, serum aliquots were stored at -80 °C.

### Glucose Tolerance Testing

At the conclusion of 4 weeks on either high glucose water or vehicle, mice were fasted overnight for 8 hours prior to glucose tolerance testing. Prior to glucose injection (time = 0 minutes), blood was drawn by tail bleed and blood glucose was measured using Contour Next Blood Glucose Monitor to obtain fasting blood glucose levels. A 20% glucose solution was prepared and autoclaved. Mice were then given an intraperitoneal (IP) injection of 2g/kg glucose. Blood glucose was measured at 15, 30, 60, 90, and 120 minutes post-injection. Mice were sacrificed using CO_2_ inhalation followed by cervical dislocation. Mice that underwent glucose tolerance testing were not utilized for downstream tissue analysis.

### Tissue Lysis and Western Blot

Liver tissues were cut on dry ice (30-40 mg sections) and homogenized in RIPA buffer (150 mM NaCl, 1% NP-40, 0.5% sodium deoxycholate, 0.1% SDS, and 50 mM Tris-HCl pH 7.4) with protease (Pierce™ Protease Inhibitor Mini Tablets, EDTA-free) and phosphatase (PhosSTOP, Roche) inhibitors. Homogenate was centrifuged (15,000 rcf, 1 hour, 4°C) and supernatant was used for immunoblotting. Lysates were frozen at -80°C overnight before protein quantification. Protein concentration was determined using BCA Assay (Pierce™ BCA Protein Assay Kit). For all protein targets, 15 µg of protein was prepared with 2-mercaptoethanol (1610710, Bio-Rad), RIPA, and LDS sample buffer (B0007, Invitrogen) according to the manufacturer’s protocols and without heating, then was loaded into a 4%–12% bis-tris 15-well gel (NW04125BOX, Invitrogen). Gels were run at 100 V for 1 hour and transferred to a low fluorescence PVDF membrane (Bio-Rad) using a Trans-Blot Turbo Transfer System (Bio-Rad). Membranes were blocked with 5% blocking grade blocker (Bio-Rad) in 1X Tris Buffered Saline with Tween 20 (TBST) (9997S, Cell Signaling Technologies), washed 3 × 5 min with TBST at room temperature, and incubated with primary antibodies in 5% BSA (BP9704-100, Fisher Scientific) overnight at 4°C. Membranes were washed 3 × 5 min with TBST and incubated in secondary antibody in 5% blocking grade blocker for 1 hour at room temperature. Membranes were washed 3 × 5 min with TBST at room temperature before imaging on a ChemiDoc MP Imager (Bio-Rad). A list of antibodies used and their dilutions is provided in Supplementary Table 1.

### Gene Expression Analysis of Liver Tissues

RNA was extracted from liver tissues (cut on dry ice into 20-25 mg sections) using the RNeasy Plus RNA isolation kit (74136, Qiagen). mRNA was quantified using a SpectraDrop Microvolume Plate on a Spectramax i3x plate reader (Molecular Devices), and 1,000 ng was added to iScript Reverse Transcription Supermix (1708841, Bio-Rad). A C1000 thermocycler (Bio-Rad) was used for reverse transcription. A total of 0.2 ng cDNA was loaded into a master mix of amplification primer and iQ SYBR green (1708882, Bio-Rad) before amplification, and gene expression was observed using a CFX Connect Real-Time PCR System (Bio-Rad). Primers used were from work by Aron *et. al*.^21^ and a list of sequences has been provided in Supplementary Table 2.

### Tissue Digestion for ICP-MS Analysis

Frozen liver tissues were cut into 30-60 mg sections on dry ice with Miltex surgical scissors that were cleaned between samples with ethanol. The sections were digested in an Eppendorf ThermoMixer Model 5382 in loosely capped 15-mL Falcon tubes (Fischer Scientific) with concentrated TraceMetal Grade (TMG) HNO_3_ (A509P500, Fisher Scientific) at 65°C for 1 hour using a constant digestion factor of 100 mg tissue/mL HNO_3_. The digested sample volumes were adjusted with 18.2 MΩ/cm MilliQ Water (MQW) to reach the original acid volume, then diluted 20-fold with MQW to reduce the acid matrix to 5% conc. TMG HNO_3_ prior to analysis by ICP-MS.

### ICP-MS Instrumentation and Procedure

Digested liver tissue samples and a digestion process water blank acidified to 5% conc. TMG HNO_3_ were analyzed by the Interdisciplinary Center for Plasma Mass Spectrometry, University of California, Davis, using an Agilent 8900 ICP-MS Triple Quadrupole instrument (Agilent Technologies) equipped with Ni interface cones and an x-Lens. The instrument was tuned and calibrated prior to sample analysis and operated in MS/MS He Mode using a 3-point peak pattern, three replicates per injection, and 50 sweeps per replicate. An Agilent SPS 4 Autosampler with a 0.5 mm ID sample probe was used to introduce samples, external calibration standards, Quality Control (QC) standards, and blanks to the ICP-MS using a 0.1 rps peristaltic pump, where they were mixed at a ∼17:1 ratio with a custom internal standard solution using a mixing tee. A 400 µL/min Micro Mist nebulizer was used to generate an aerosol in a 2°C temperature controlled double pass Scott-type spray chamber leading to a 1550 W plasma. He was used as a collision cell gas to manage polyatomic interferences.

### Standard Preparation and ICP-MS Calibration

Single-element Fe, Cu, and Zn calibration standards, QC standards, and a custom internal standard solution with Sc, Ge, Y, In, and Bi were prepared from single-element standards (Inorganic Ventures). All solutions were prepared with conc. TMG HNO_3_ and MQW with all final solutions at 5% conc. TMG HNO_3_. A calibration blank and seven calibration standards with concentrations from 5 to 1000 ppb were prepared and analyzed to establish linear calibration equations and determine detection limits for Fe, Cu, and Zn using Ge as the internal standard.

### Quality Assurance and ICP-MS Data Processing

Quality assurance was established by analyzing and evaluating a digestion process water blank, an independent source NIST 1643f – Trace Elements in Water standard and blank for initial calibration and blank verification, and QC standards and blanks before and after no more than ten sample injections for continuing calibration and blank verification and to monitor instrument drift. Raw data was processed with MassHunter 4.6 ICP-MS software (G7201C, Version C.01.06, Agilent) to determine Fe concentrations in the digested liver samples as ng Fe/mg liver.

### Enzyme-Linked Immunosorbent Assays (ELISA) of Serum Iron Proteins

Serum iron markers were all quantified using enzyme-linked immunosorbent assays from Abcam. Serum was diluted 1:100,000 for serum transferrin measurement and run according to manufacturer’s instructions (ab157724, Abcam). For serum ferritin and serum transferrin receptor assays serum was diluted 1:40 and assays were run according to manufacturer recommendation (ab157713, ab243674, Abcam). Final absorbance measurements for all assays were performed on Spectramax i3x plate reader (Molecular Devices).

### Preparation of Holo-transferrin and Holo-ferritin LC-ICP-MS Calibration Curve

All LC-ICP-MS methods are modified from a previously published study by Neu *et al*^22^. Crystallized human holo-transferrin (Sigma Aldrich) and crystallized holo-ferritin from equine spleen (Millipore Sigma) were used to prepare a calibration curve containing both standards. A stock solution of transferrin was prepared by dissolving 0.00351 g of holo-transferrin in 0.975 mL of 10 mM Tris buffer to afford a 5000 ppb solution. A stock solution of ferritin was prepared by diluting 100 µL of holoferritin in 900 µL of 10 mM Tris buffer to afford a 1,000,000 ppb solution. A stock solution containing holo-transferrin and holo-ferritin (Stock 1) was prepared by adding 200 µL of the transferrin stock solution and 10 µL of the ferritin stock solution to 790 µL of 10 mM Tris buffer. The calibration curve consisting of eight points ranging from 50-1000 ppb transferrin and 500-10000 ppb ferritin was prepared via serial dilution of Stock 1 using 10 mM Tris buffer and analyzed to establish linear calibration equations and determine detection limits for transferrin and ferritin by measuring the time resolved Fe signal using Ge as the internal standard. All solutions containing holo-transferrin and holo-ferritin were kept on ice until use.

### Serum Preparation for LC-ICP-MS

For LC-ICP-MS experiments, serum samples were thawed on ice and prepared by gently agitating prior to adding 20 µL of serum to 80 µL of 10 mM Tris buffer, pH 7.4. All samples containing mouse serum were stored in ice until use.

### Liquid Chromatography-ICP-MS Analysis of Serum Proteins

Standard solutions and serum samples were injected at room temperature onto an Agilent 1260 Infinity II Quaternary Pump controlled by Agilent Lab Advisor software (Version 2.20.543 – Basic, Agilent) and connected to the ICP-MS at a flow rate of 0.400 mL/min using 10 mM Tris base pH 7.4 as the mobile phase. The injection volume was 20 µL and the total run time was 18 min. Separation was achieved by size exclusion chromatography with an Agilent Bio SEC-3 column (300A, 4.6 × 300 mm, 3 µm) connected to a Bio SEC-3 guard column (300A, 4.6 × 50 mm, 3 µm). The ICP-MS was operated using time resolved analysis in single quad mode using He as the collision cell gas. Point-to-point count correction was used with Ge set as the online internal standard. The Agile2 integrator was used for peak integration. Blanks were analyzed after the calibration standards, after the initial set of six serum samples, and after the final set of six serum samples followed by a calibration verification standard for blank verification and to monitor instrument drift. Raw data was processed with MassHunter 4.6 ICP-MS software and the optional Chromatographic Analysis software (G7201C, Version C.01.06, Agilent) to determine transferrin and ferritin concentrations in the mouse serum samples.

### Cell Stimulation and Western Blot

HepG2 cells were seeded in a clear-bottom 6-well plate at 200,000 cells per well and allowed to adhere overnight in complete DMEM media. Cells were then left to incubate for 48 hours in low glucose (1 g/L) or high glucose (4.5 g/L) complete DMEM, with and without ferrous ammonium sulfate. Ferrous ammonium sulfate stock was prepared fresh in nanopure water (Millipore) at a concentration of 2.5 mM and diluted to 10 µM in media. After 48 hours, cells were treated with either metformin or deferoxamine (DFO). Metformin stocks were prepared in nanopure water at a concentration of 200 mM and diluted to 2 mM in media. DFO stocks were prepared in nanopure water at a concentration of 250 µM and diluted to 25 µM in media. Cells were incubated in either metformin or DFO for 24 hours prior to harvest. Cells were lysed in RIPA buffer (150 mM NaCl, 1% NP-40, 0.5% sodium deoxycholate, 0.1% SDS, and 50 mM Tris–Cl pH 7.4) with an EDTA-free protease inhibitor (PIA32955, Thermo Fisher Scientific) and a phosphatase inhibitor (4906845001, MilliporeSigma). Lysates were vortexed and allowed to sit on ice for 10 min prior to being cleared by centrifugation at 15,000 rcf at 4°C. Lysates were frozen at −20°C prior to protein quantification using BCA assay (71285-3, Invitrogen). For all protein targets, 10 µg protein was prepared with 2-mercaptoethanol (1610710, Bio-Rad), RIPA buffer, and LDS sample buffer (B0007, Invitrogen) according to the manufacturer’s protocols and without heating, then loaded into a 4%–12% bis–tris 15-well gel (NW04125BOX, Invitrogen) for probing all proteins. Gels were run and western blotting was performed according to protocols described above.

## RESULTS AND DISCUSSION

### Study Design

This study is intended to focus on hyperglycemic conditions. To determine the effect of high sugar diet on metabolic and iron regulation, C57BL/6J mice were given 10% glucose supplemented water (high glucose drink, HGD) for 28 days as glucose is the primary energetic fuel in mammals. Previous studies have used between 10 and 30 percent dietary sugar supplemented water^8,23–29^. Our study focuses solely on glucose supplementation due to fructose’s dual role as both a nutrient and a metabolic signal for sugar intake^30^. Using combinations of sugars introduces confounding effects from this multifunctional role. In order to evaluate whether a four-week course of HGD was sufficient to cause metabolic changes, mice were weighed weekly and both serum and hepatic metabolic markers were measured. As the liver is both a major metabolic and iron regulator systemically, our studies will focus on both hepatic metabolic and iron status.

### High Glucose Supplementation Causes Weight Gain and Increased Fat Production

Metabolic markers were screened in order to determine the phenotype of mice on HGD. Mice given the HGD showed increased weight gain starting seven days after beginning high glucose diet. At the end of the four-week period, mice on HGD exhibited a 20 percent increase in weight. Comparatively, mice on non-supplemented water exhibited less than 10 percent weight gain during the same four-week period (Figure 1B, 1C). Differences in weight gain may be attributed to increased peripheral fat production. Mice on HGD had significantly more white adipose tissue than vehicle mice (Figure 1D). At the end of the 4-week period, fasting blood glucose levels were analyzed and intraperitoneal glucose tolerance testing (IPGTT) was performed. These parameters inform on the insulin-mediated uptake of glucose from the blood stream. Fasting blood glucose levels of mice on HGD were higher, though not significantly, compared to vehicle mice (Figure 1G). Upon i.p. injection of glucose, the mice on HGD showed an initial elevation of blood glucose to a level modestly higher than the vehicle mice; but the blood glucose of the two groups lowered to the same levels 120 minutes after glucose injection (Figure 1E). These results show that while a 4-week exposure to HGD does lead to increased weight gain, fat levels, and fasting blood glucose levels, it does not impair glucose tolerance. Similar results have been previously observed in mice given 15% fructose over a 4-week time period^28^ in contrast to the glucose intolerance observed with longer exposures^24^. It is thus likely that the 4-week exposure is an insufficient duration to affect glucose tolerance.

**Figure 1:**
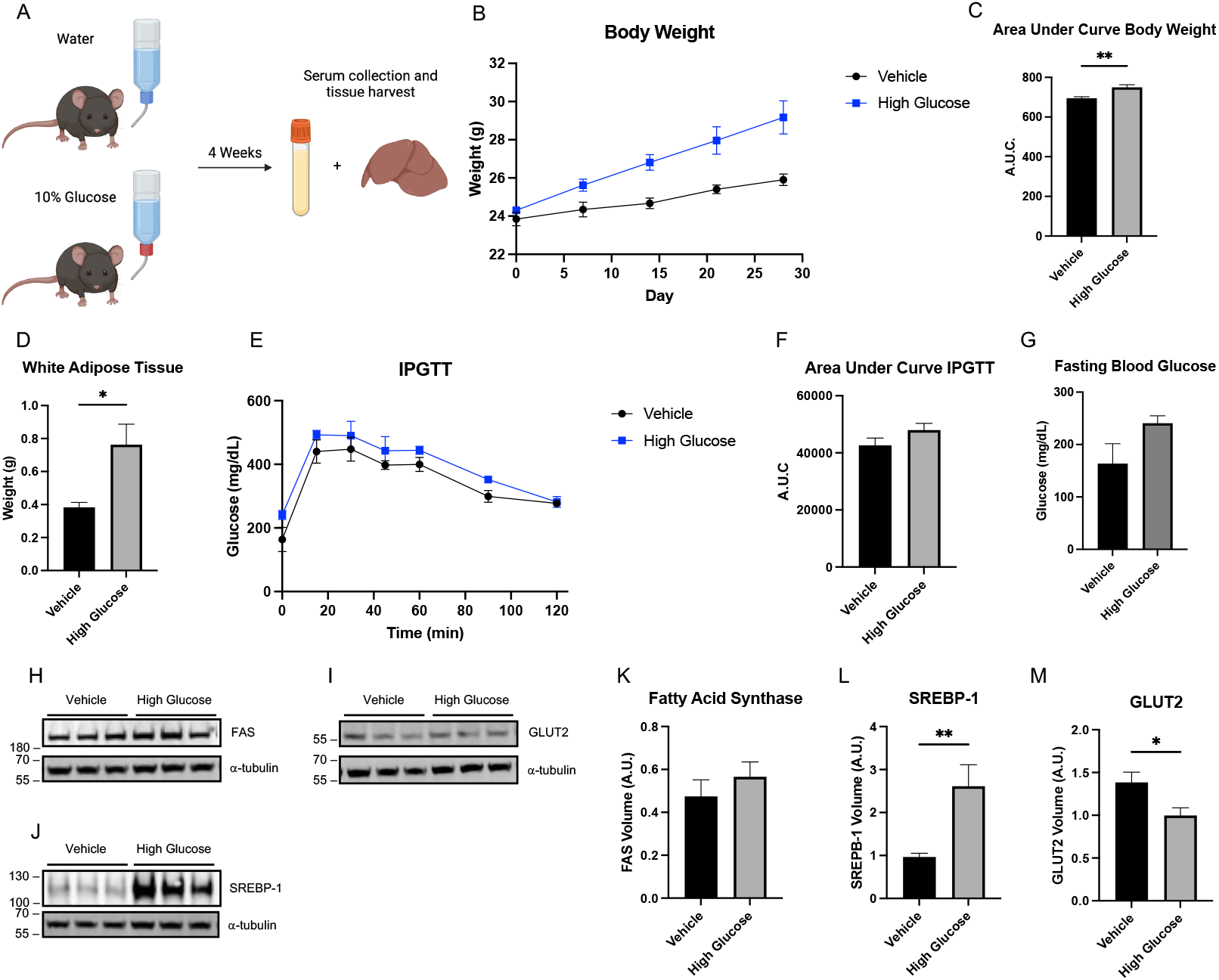
HGD supplementation induces hyperglycemia and stimulates fat production. **A** Schematic representation of experimental design for of high glucose drink supplementation in C57BL/6J mice. **B** Body weight of HGD and vehicle groups over time, measured weekly (n=6). **C** Area under the curve of body weight of HGD and vehicle groups. **D** Weight of white adipose tissue collected after sacrificing HGD and vehicle mice (n=6). Tissues were weighed and collected on day 28. **E** IPGTT was performed on mice given HGD and vehicle on day 28. Blood glucose was measured at 0, 15, 30, 45, 60, 90, and 120 minutes (n=3). **F** Area under the curve for IPGTT. **G** Fasting blood glucose of HGD and vehicle mice (t=0 of IPGTT). **H** Representative western blot of fatty acid synthase (FAS) in liver lysates of HGD and vehicle groups. **I** Representative western blot of glucose transporter 2 (GLUT2) in liver lysates of HGD and vehicle groups. **J** Representative western blot of sterol regulatory binding protein 1 (SREBP-1) in liver lysates of HGD and vehicle groups. **K** Densitometry analysis of FAS western blot for HGD and vehicle groups (n=6). **L** Densitometry analysis of SREBP-1 western blot for HGD and vehicle groups (n=6). **M** Densitometry analysis of GLUT2 western blot for HGD and vehicle groups (n=6). All data with error bars are displayed as Mean ± SEM. α-tubulin was utilized as a loading control for all western blot analysis. One-tailed t-test was used for all statistical analyses. *p-value <0.05, **p-value < 0.01.

Protein expression levels of metabolic regulators were analyzed using western blot. Fatty acid synthase (FAS) is an enzyme catalyzing de novo synthesis of fatty acids^31^. In the liver, FAS has been shown to shift toward energy storage via fat production when there is an excess of nutrients^31^. Mice on HGD show a modest, though not statistically significant, increase in the expression of FAS in the liver (Figure 1K). Sterol regulatory element binding proteins (SREBPs) regulate lipid homeostasis. In the liver, heightened expression of SREBP-1 can activate pathways for the biosynthesis of both fatty acid and cholesterol^32^. Mice on HGD show a 2.5-fold increase in liver SREBP-1 levels compared to the vehicle group (Figure 1L). A prior study with sucrose supplementation also showed increases in SREBP-1 in the liver of rats^26^. However, despite the elevation in both FAS and SREBP-1 expression, Oil O red staining did not show an increase fat macrodroplets in hepatic tissues (Figure S1). Additionally, liver tissue weights were not significantly different between the high glucose and vehicle groups (Figure S2). As with the maintenance of a glucose-tolerant state, the lack of increase in fat macrodroplets may indicate that the 4-week mark precedes fat accumulation.

Interestingly, mice on HGD showed reduced expression of glucose transporters 2 (GLUT2) (Figure 1M). GLUT2 is essential for hepatic glucose absorption, accounting for over 97% of all glucose transporters in hepatocytes^33^. The observed decrease in GLUT2 expression may be indicative of hepatic glucose accumulation due to the high glucose supplementation. A similar reduction in *Glut2* mRNA levels were observed in a study of mice given 10% high fructose corn syrup (HFCS) in water for 12 weeks^24^. Since HFCS is composed of 55% fructose and 45% glucose, the comparable effects on glucose metabolism are not surprising. The western blot data, combined with the observed metabolic phenotypes, indicate that 4 weeks of HGD exposure induces a hyperglycemic state without causing glucose intolerance. This exposure also enhances hepatic fat production, shifting metabolism toward energy storage.

### HGD Intake Increases Hepatic Iron Uptake and Accumulation

Previous research investigating the relationship between iron and glucose regulation has focused on iron as a driver of type II diabetes or chelation as a treatment strategy. Consequently, the majority of studies to date have used either knockout models of hemochromatosis or diabetic (both genetic and diet-based) rodent models^35–39^. However, the specific impacts of high sugar diets on iron regulation and trafficking are understudied ^14,34^. Understanding the directionality of which hyperglycemia and iron regulation are related may help reveal alternate therapeutic targets in treating metabolic dysfunction. To address this gap, we sought to characterize the iron status of mice given HGD. Given the central role of liver in iron trafficking, we assessed hepatic iron status with respect to both total iron levels and iron regulatory proteins. Total iron levels in liver tissue were quantified using inductively coupled plasma mass spectrometry. Mice on HGD exhibited a 30% increase in total liver iron levels compared to the vehicle group (Figure 2A). This increase is particularly notable as both HGD and vehicle mice are fed the same chow and therefore have no difference in the iron content of either their drink or diet. Increases in hepatic iron levels were accompanied by changes in expression levels of iron regulatory proteins, as measured by real-time quantitative reverse transcription PCR (RT-qPCR) and western blot analysis. Western blot analysis of TFRC, the transferrin receptor, shows a two-fold increase with the HGD supplementation when compared to mice given the vehicle (Figure 2G). Transferrin is the protein responsible for transporting iron in the serum. Transferrin receptors on cell surfaces bind transferrin and initiate endocytosis of the protein to release and internalize iron into the cell^40^. An increase in transferrin receptor expression is indicative of higher levels of transferrin-bound iron uptake. This is commonly seen in cancer cells, which enhance iron uptake to support cell proliferation. However, changes in iron homeostasis in non-cancerous cells have also been shown to affect TFRC expression^41,42^. Additionally, TFRC has been implicated in influencing systemic iron status by regulating hepcidin expression^43^. Increased expression of ferritin heavy chain (FTH1) was also seen in mice given HGD (Figure 2H). Similarly to TFRC, ferritin expression is increased in hepatocytes due to iron overload^42^ (Figure 2J). Previous studies utilizing cell models have found that sugars including fructose, sucrose, and high fructose corn syrup increase iron bioavailability and increase intracellular ferritin levels^44^. This is thought to be due to sugars either having chelation effects or altering oxidation state of iron thus making iron more bioavailable and leading to increased iron levels. Glucose, however, has not been previously reported to increase iron storage. Additionally, the mechanism by which these sugars increase iron uptake remains to be elucidated. Iron accumulation appears to be limited to transferrin-bound iron as there is a decrease in divalent metal transporter 1 (DMT1) expression (Figure 2E) which is a metal transporter implicated in non-transferrin-bound iron uptake. However, alterations in DMT1 expression may have no distinct correlation with iron accumulation as studies suggest the protein to be unnecessary for sufficient hepatic iron uptake^45^. In addition to the transporters and storage proteins, we investigated effects on the hepcidin-ferroportin axis. The liver produces hepcidin in response to systemic increases in iron to effectively reduce iron absorption from the intestines. When hepcidin binds to ferroportin on duodenol enterocytes, it triggers the degradation of FPN1, halting iron export from the duodenum and into the plasma^46^. RT-qPCR analysis showed an increase in the mRNA expression of *hamp* and *fpn1* genes (which encode for the hepcidin and ferroportin, respectively) in the HGD group relative to the vehicle group (Figure 2B). Interestingly, analogous changes in protein expression for ferroportin were not observed. The elevation of *hamp* may indicate that the HGD shifts the mouse livers to decreased intestinal iron absorption. Overexpression of the *hamp* gene in mouse liver has previously been found to occur during iron overload^47^. However, the coinciding increase in *fpn1* is surprising, and may indicate a counterbalance to regulate hepatic iron levels. Indeed, elevated *fpn1* expression has previously been observed in metabolic disorders like metabolically associated fatty liver disease (MAFLD)^48^. Taken together, these results suggest that the metabolic state caused by HGD leads to increased uptake of transferrin-bound iron resulting in hepatic iron overload.

**Figure 2:**
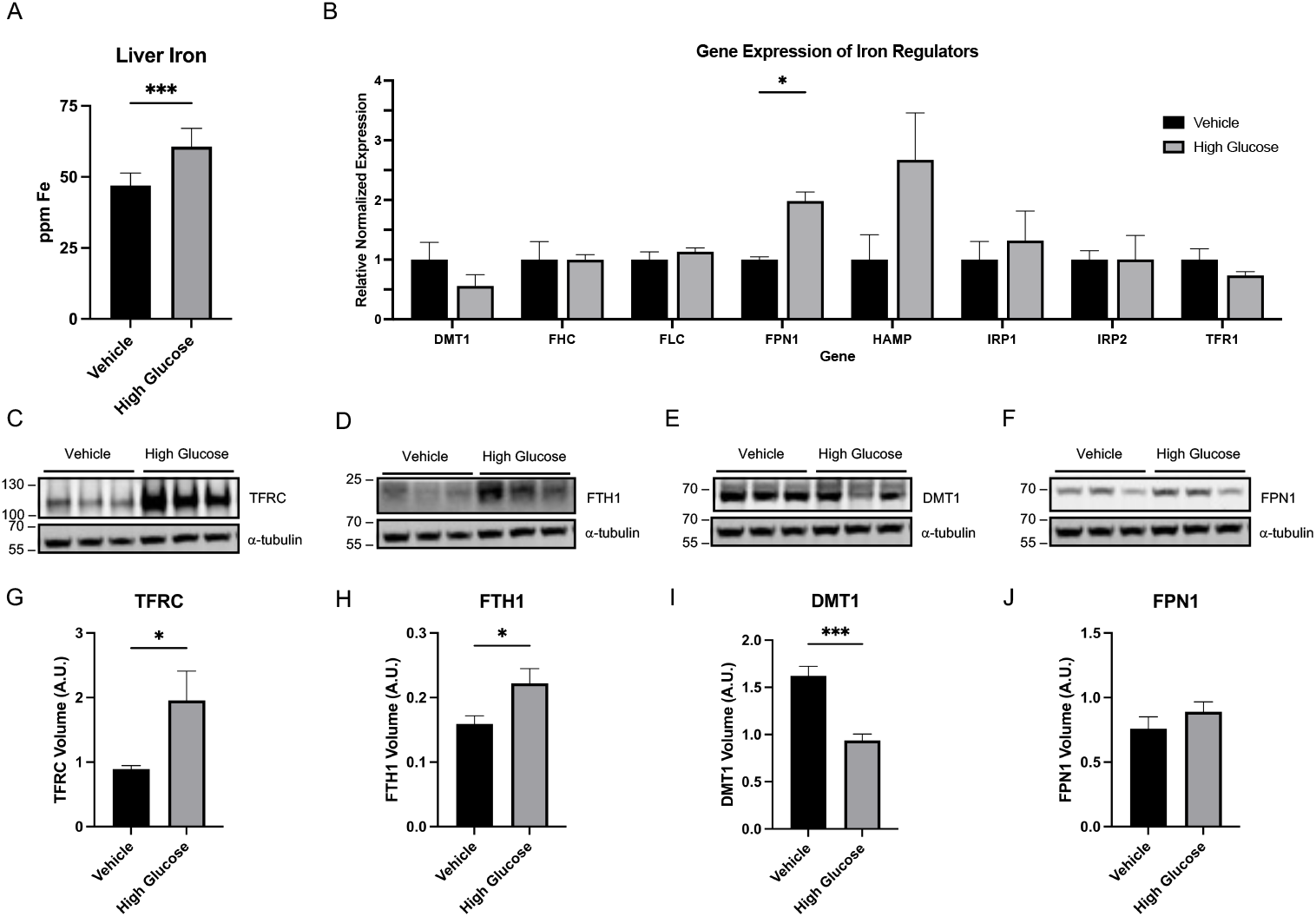
HGD induces hepatic iron overload. **A** Inductively coupled plasma mass spectrometry (ICP-MS) analysis of liver tissue of HGD and vehicle mice. Samples were normalized by tissue weight. (n=6) **B** Real-time quantitative PCR of iron regulators in the liver of HGD and vehicle mice. Gene expression analysis was performed for divalent metal transporter 1 (DMT1), ferroportin (FPN1), transferrin receptor 1 (TFR1), ferritin heavy chain (FHC), ferritin light chain (FLC), hepcidin (HAMP), iron-responsive element-binding protein 1 (IRP1), and iron-responsive element-binding protein 2 (IRP2). (n=6) **C** Representative western blot of transferrin receptor (TFR1) in the liver of HGD and vehicle mice. **D** Representative western blot of ferritin heavy chain 1 (FTH1) in the liver of HGD and vehicle mice. **E** Representative western blot of divalent metal transporter 1 (DMT1) in the liver of HGD and vehicle mice. **F** Representative western blot of ferroportin (FPN1) in the liver of HGD and vehicle mice. **G** Densitometry analysis of TFRC western blot for HGD and vehicle mice (n=6). **H** Densitometry analysis of FTH1 western blot for HGD and vehicle mice (n=6). **I** Densitometry analysis of DMT1 western blot for HGD and vehicle mice (n=6). **J** Densitometry analysis of FPN1 western blot for HGD and vehicle mice (n=6). Data displayed as Mean ± SEM. α-tubulin was utilized as a loading control for all western blot analysis. One-tailed t-test was used for all statistical analyses. *p-value < 0.05, **p-value < 0.01.

### Hepatic Iron Overload is Induced by Increased Uptake of Transferrin-Bound Iron

The observed changes in iron markers in the liver in response to the HGD administration point to either systemic increases in iron levels or tissue specific hepatic iron overload. Serum iron markers were thus evaluated to further untangle the HGD-dependent effects on iron status. Specifically, we sought to interrogate whether hepatic iron overload was a manifestation of systemic overload due to increased dietary iron absorption or a rearrangement of systemic iron toward hepatic accumulation.

We first determined whether changes were occurring in the quantities of the established serum iron proteins by ELISA. The measurements showed no differences in the serum levels of either ferritin or transferrin with HGD supplementation (Figure 3A, Figure S3). Interestingly, serum transferrin (STfR) exhibits a 30 percent increase in serum levels in the HGD group relative to vehicle (Figure 3B). STfR is known to correlate with total body TfR concentration^49^. This correlation is present in our samples, as elevated TFRC expression in the liver lysates are also observed in the HGD group (Figure 2C, 2G).

**Figure 3:**
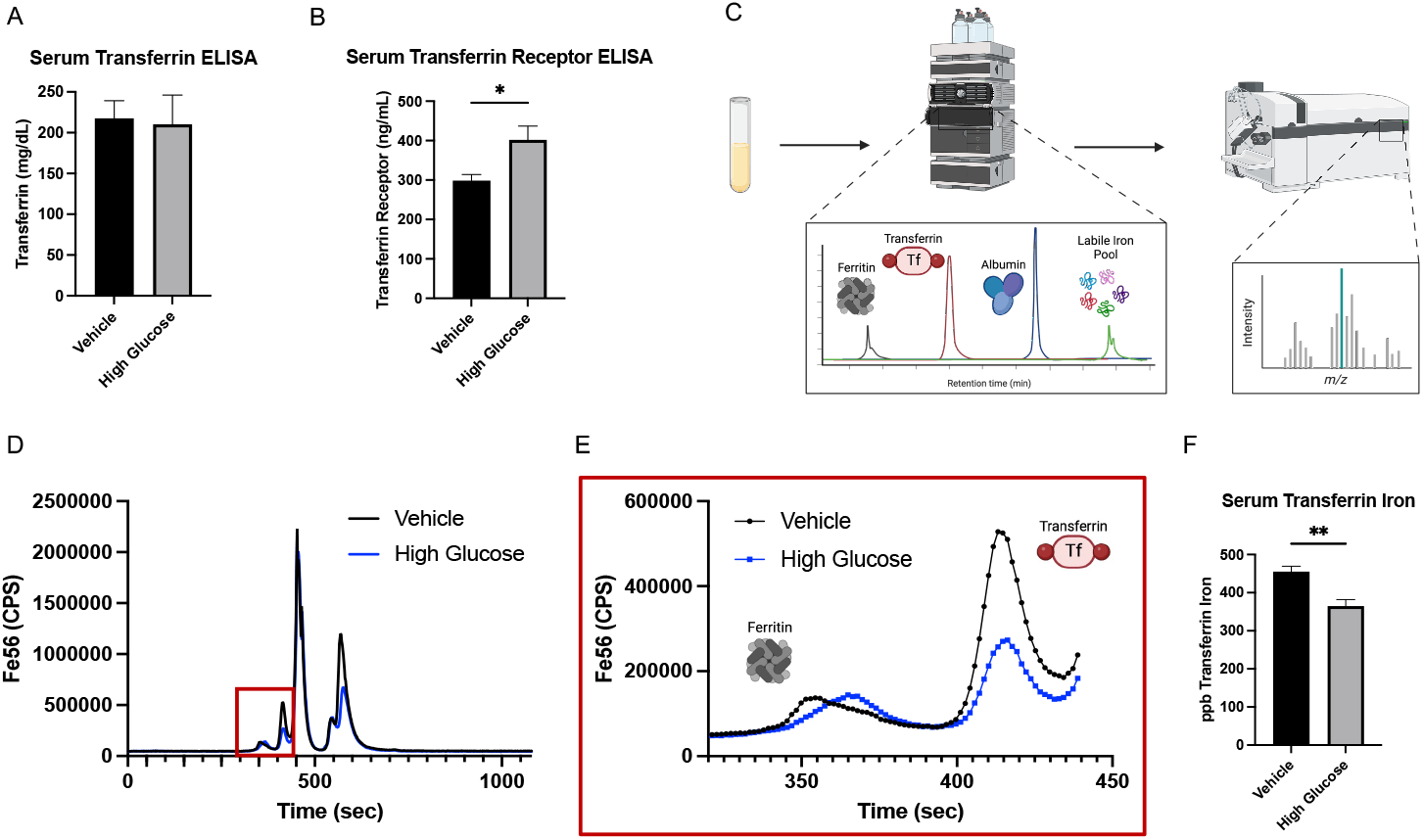
HGD increases transferrin-bound iron uptake. **A** Enzyme-linked immunosorbent assay (ELISA) was used to quantify serum transferrin levels in HGD and vehicle mice (n=6). **B** ELISA quantification of transferrin receptor in serum of HGD and vehicle mice (n=6). **C** Schematic of liquid chromatography inductively coupled plasma mass spectrometry (LC-ICP-MS) used to quantify iron content of serum proteins. **D** Representative spectra of the serum of HGD and vehicle mice measured using (LC-ICP-MS). Peak differences at 650 seconds due to differences in hemolysis of samples. **E** Magnification of ferritin and transferrin peaks in representative spectra of serum LC-ICP-MS of HGD and vehicle mice. Differences in peak shape of ferritin may be due to different heavy and light chain compositions of ferritin. **F** Serum transferrin iron levels were quantified via standard curve and peak integration of LC-ICP-MS runs. n=6 mice were quantified per condition. Data displayed as Mean ± SEM. One-tailed t-test was used for all statistical analyses. *p-value < 0.05, **p-value < 0.01.

ELISA measures total protein levels but does not capture the metalation state of metalloproteins. Shifts in metal status can influence metal-loading while keep protein levels unchanged, a phenomenon particularly relevant in the context iron balance. Ferritin, for example, adjusts its iron loading to both store and release iron in response to biological demands, independent of changes to its protein expression^50,51^. This concept is also recognized in clinical practice with transferrin, where transferrin saturation with iron serves as an independent measure of iron status, separate from transferrin protein levels^52^. Iron loading independent of protein levels is also intrinsic to iron uptake pathways: iron-bound (or holo) transferrin binds to the transferrin receptor to mediate endocytosis; then, acidification of endosomes liberates the bound iron for cellular use, while apo-transferrin is recycled and returned to circulation, effectively trafficking iron without altering protein levels^53^. Given these mechanisms, we further investigated serum iron status by assessing the iron loading of the serum proteins.

We assessed the iron loading of serum transferrin and ferritin via liquid chromatography-inductively coupled plasma-mass spectrometry (LC-ICP-MS). In this experiment, serum components are fractionated via HPLC on a size exclusion column then directly flowed into an ICP-MS detector for iron quantification of the fractions. When used alongside standards, retention times can be used to benchmark the species to known iron-containing proteins, allowing for iron distribution and loading to be measured (Figure 3C). Sera from both HGD and vehicle mice were analyzed by this method. (Figure 3D, 3E). Iron levels within serum ferritin showed high variability independent of drink content (Figure S4). Transferrin, however, showed a 20 percent decrease in iron levels in the HGD group relative to vehicle (Figure 3F). As protein levels of transferrin were previously determined to be equal between both groups, this points to a decrease in transferrin saturation in HGD group. This observation taken alongside the increased expression of TFRC in the liver, increased STfR, and increased hepatic iron levels suggests that HGD may stimulate an increase in uptake of transferrin-bound iron in the liver resulting in hepatic iron overload (Figure 4). The underlying driving force for the increase in hepatic iron levels remains unclear, however, it may be that increased fat synthesis driven by HGD is an iron dependent process thus leading to accumulation of iron in the liver. The relationship between iron accumulation and fat production has also been seen in diseases such as MAFLD^17,54^. While further studies are necessary to fully elucidate the mechanism linking the two, this phenomenon highlights the potential for iron as a therapeutic target for metabolic diseases.

**Figure 4:**
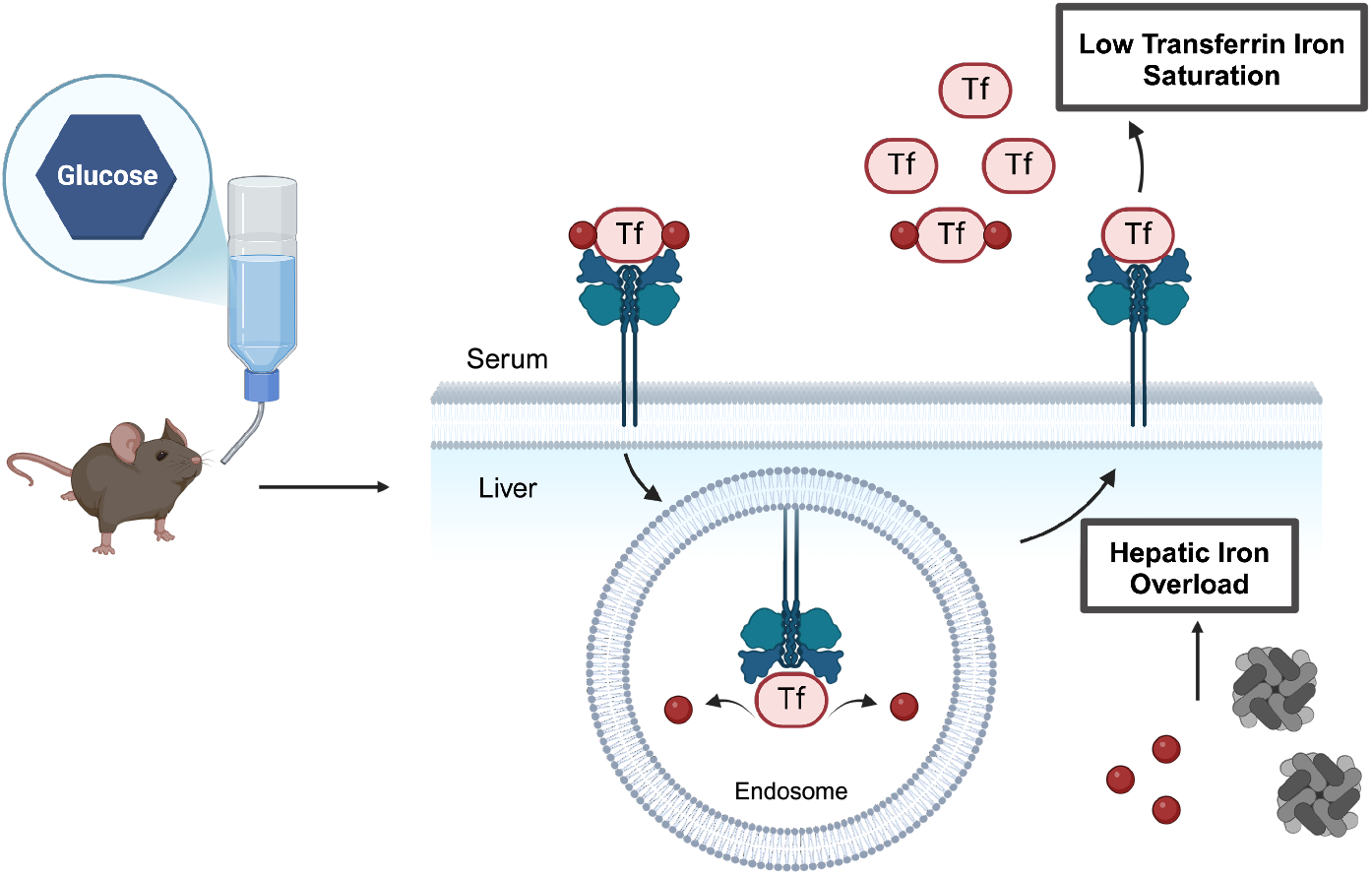
HGD shifts metabolism towards fat production and induces hepatic iron overload via increased uptake of transferrin-bound iron. Summary figure of iron regulation in mice given a high glucose drink. Mice show increases in liver iron stores both as increased ferritin and increased total iron levels. Serum analysis shows an increased uptake of transferrin-bound iron as the mechanism by which hepatic iron overload.

### Metformin Restores Metabolic and Iron Homeostasis in HepG2 Cells

The perturbations in iron metabolism we observed following a short-term HGD prompted us to explore whether restoration of metabolic regulation could reverse hepatic iron overload or if iron chelation could restore metabolic function. We opted to use a cell-based model with the HepG2 cell line, a human liver cell line, (Figure 5A), as it is well-studied both with respect to glucose regulation and iron chelation^55,56^. We found that exposing the cells to a combination of high glucose and high iron in the media could stimulate HepG2 cells into a state that mimics the hepatic iron overload state observed in the mice given HGD, including the increased expression of SREBP-1 (Figure 5C) and the large increase in ferritin expression (Figure 5F). To determine the effects of iron chelation on metabolic regulation, cells were treated with deferoxamine (DFO) following the high glucose and high iron exposure. DFO is an FDA approved iron chelator for iron overload disorders. It does not possess membrane permeability but chelation of extracellular iron with DFO has been shown to decrease intracellular iron stores^57^. We also investigated the effects of metformin (*N,N*-dimethylbiguanide), the most commonly prescribed anti-diabetic drug, to determine whether metabolic shift in turn could restore iron regulation. It is important to note, however, the understanding of metformin’s mechanism of action remains incomplete. Previous studies have shown that metformin can bind bioavailable copper, suggesting that its effects are at least partially driven by copper binding and sequestration^58,59^. While the effect of metformin on iron regulatory pathways is unknown, recent studies have shown that metformin could inhibit ferroptosis, an iron-associated form of cell death, in non-alcoholic fatty liver disease^60,61^.

**Figure 5:**
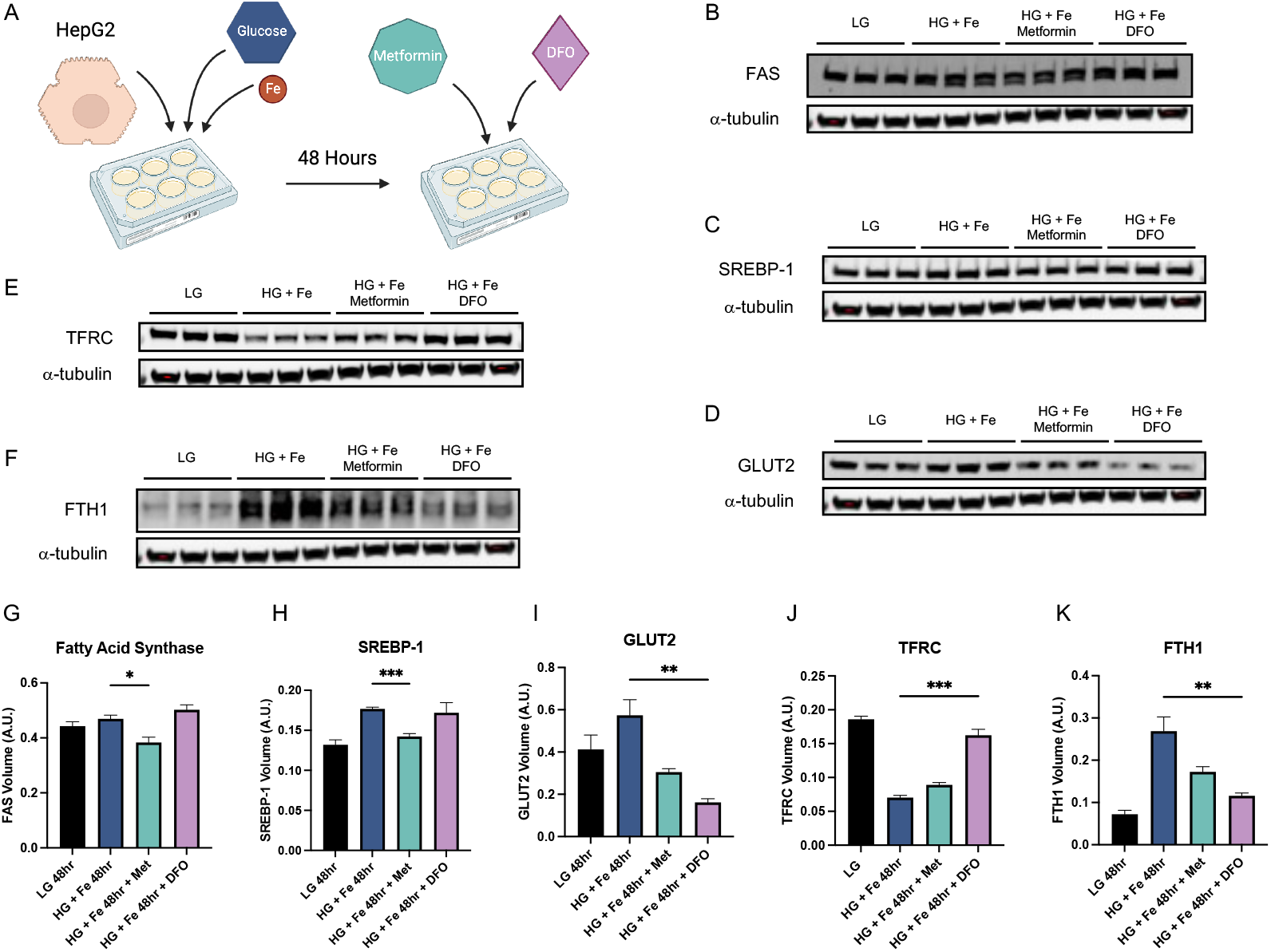
Application of a cell-based model to assess the metformin and iron chelation treatment on iron misregulation and glucose dyshomeostasis. **A** Schematic of cell stimulations. Cells were grown for 48 hours in either low glucose (1 g/L) or high glucose (4.5 g/L) high iron media (additional 10 µM). Cells were then treated with either metformin (2 mM) (green octagon) or deferoxamine (25 µM) (purple diamond), an iron chelator. **B-F** Representative western blot of cell stimulations for fatty acid synthase (FAS) (**B**), sterol regulatory binding protein (SREBP-1) (**C**), glucose transporter 2 (GLUT2) (**D**), transferrin receptor (TFR1) (**E**), and ferritin heavy chain (FTH1) (**F**). **G-K** Densitometry analysis of western blot for FAS (**G**), SREBP-1 (**H**), GLUT2 (**I**), CD71 (**J**), and FTH1 (**K**) in stimulated cells (n=3). Data displayed as Mean ± SEM. α-tubulin was utilized as a loading control for all western blot analyses. One-tailed t-test was used for all statistical analysis. *p-value < 0.05, **p-value < 0.01.

As expected, chelation with DFO restores normal TFRC and FTH1 expression levels in the high glucose/high iron-exposed cells (Figure 5E, 5F, 5J, 5K). Interestingly, chelation with DFO does not change expression of either FAS or SREBP-1 (Figure 5C, 5G), but shows a large decrease in GLUT2 expression (Figure 5B, 5G). This suggests that iron chelation is tied directly to glucose uptake. Changes in fat production markers may require a longer time scale of decreased glucose uptake than what we probed and longer treatment duration would likely result in decreased fat production. When the high glucose/high iron-exposed HepG2 cells are treated with metformin, we observe a decrease in expression of both FAS and SREBP-1 back to normal levels (Figure 5B, 5C, 5G, 5H), which is not surprising, given the known ability of metformin to affect cellular metabolism. Altering glucose metabolism is thought to be one of the primary mechanisms by which metformin treats hyperglycemia and type II diabetes^58^, and as expected, metformin decreases GLUT2 levels. Interestingly, however, the decrease in GLUT2 was not as pronounced as when the cells are treated with DFO, highlighting the potential of chelation strategies as therapeutic options in hyperglycemic conditions. In addition to the expected metabolic changes, we found that metformin treatment also influenced iron regulation. While metformin treatment did not restore TFRC levels to normal, there was a slight increase in expression, and FTH1 levels were significantly decreased with treatment. This suggests that metformin may restore normal iron regulation and reverse the cellular iron accumulation observed under high glucose and iron conditions. While there is no established precedent for iron regulation restoration as a mechanism of metformin, the involvement of mitochondria in its action, coupled with these findings, indicates that metformin might function, at least partially, by reducing intracellular iron accumulation. The restoration of metabolic function through treatment of iron misregulation with both metformin and DFO that are concomitant with changes in iron-associated proteins emphasizes the importance of understanding iron regulation in conditions that alter metabolic states.

## CONCLUSION

In this work, we explore the mechanism of iron regulation in the context of hyperglycemic states. The Western diet has led to an increased incidence of metabolic disorders, yet the mechanisms that drive disease progression remain poorly understood. Without detailed mechanistic understanding of disease onset, treatments remain limited and existing drugs lack well established mechanisms of action. Our findings show that a high glucose drink can shift metabolic regulation and lead to alterations in systemic iron homeostasis. Hepatic iron accumulation is driven by an increase in uptake of transferrin-bound iron despite no changes to dietary iron intake. By applying clinically used agents to a cellular model that mimics the hyperglycemic and iron overload conditions seen in our mouse model, we found that the glucose regulatory drug metformin is able to restore iron regulation by decreasing excess intracellular iron, while the iron chelator DFO can significantly alter glucose transport. These observations provide new mechanistic insights into drug action that link iron homeostasis to glucose balance. Broadly, our study underscores the importance of investigating molecular basis of shifts in metal homeostasis in metabolic diseases, and how such research could reveal new druggable targets.

