## Supplemental Info for "High Glucose Diet Induces Hepatic Iron Overload Contributing to Metabolic Dysfunction"

**SUPPORTING TEXT**

**Tissue Sectioning.** Livers were sectioned at -20 ºC on a Thermo Fisher Scientific CryoStar

NX70 Cryostat (Thermo Fisher Scientific). Sections were cut at a thickenes of 10 µm and thaw mounted onto Fisherbrand™ Premium Plain Glass Microscope Slides (Thermo Fisher Scientific).

**Oil O Red Staining.** Prior to staining, mounted tissues were fixed in 4% paraformaldehyde in PBS (J61899.AK, Thermo Fisher Scientific) for 15 minutes. Slides were then rinsed in water. A 0.5% stock solution of Oil Red O (ORO) (A12989.22, Thermo Fisher Scientific) was prepared by dissolving 0.5 g of ORO with 100 mL of 100% isopropanol (IPA) (447080010, Thermo Fisher Scientific). Solution was heated in a water bath at 65 ºC for until fully dissolved. A working solution was then prepared by diluting 30 mL of stock with 20 mL of nanopure water. Solution was filtered using filter paper and a ceramic funnel three times to remove excess stain. Fixed slides were placed in 60% IPA for 2 minutes and then placed in ORO working solution for 15 minutes. After staining, slides were rinsed in water for 3 minutes. 80% glycerol in water was used as mounting medium. Slides were mounted and imaged using an EVOS FL microscope (Invitrogen) with 20x magnification.

**SUPPORTING FIGURES**

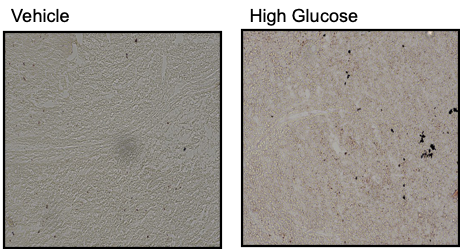

**Figures S1.** Oil O red staining was performed on cryosections of liver tissues for both HGD and vehicle mice. Differences in hepatic fact accumulation between conditions were not observed.

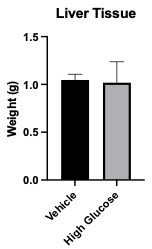

**Figure S2.** Weight of liver tissue collected of HGD and vehicle mice (n=6). Tissues were harvested and weighed after sacrificing the mice on day 28. Data displayed as Mean ± SEM.

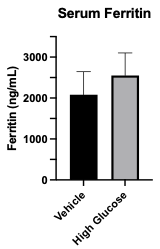

**Figure S3.** Enzyme-linked immunosorbent assay (ELISA) was used to quantify serum ferritin levels in HGD and vehicle mice (n=6). Data displayed as Mean ± SEM.

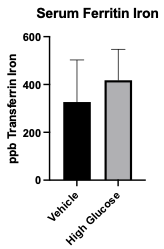

**Figure S4.** Serum ferritin iron levels were quantified via standard curve and peak integration of LC-ICP-MS runs. n=6 mice were quantified per condition. Data displayed as Mean ± SEM.

**SUPPORTING TABLES**

**Table S1.** Antibodies used for western blot and their dilutions. Primary antibodies were diluted in 5% BSA containing 0.05% sodium azide. Secondary antibodies were diluted in 5% blotting grade blocker.

| **Primary Antibodies** | | | |
| --- | --- | --- | --- |
| **Name** | **Dilution** | **Manufacturer** | **Catalogue #** |
| FAS | 1:2000 | Cell Signaling Technology | 3180S |
| GLUT2 | 1:2000 | EMD Millipore Corp | 07-1402-I |
| SREBP-1 | 1:2000 | Santa Cruz Biotechnology | sc-365513 |
| α - tubulin | 1:5000 | Invitrogen | MA1-80017 |
| TFRC | 1:2000 | Santa Cruz Biotechnology | sc-52256 |
| FTH1 | 1:1000 | Santa Cruz Biotechnology | sc-376594 |
| DMT1 | 1:2000 | Santa Cruz Biotechnology | sc-166884 |
| FPN1 | 1:2000 | Novus Biologicals | NBP1-21502 |
| **Secondary Antibodies** | | | |
| **Name** | **Dilution** | **Manufacturer** | **Catalogue #** |
| AF-800 | 1:2500 | Invitrogen | A32789 |
| AF-555 | 1:2500 | Invitrogen | A21428 |
| AF-647 | 1:2500 | Invitrogen | A21448 |

**Table S2.** Sequences of primers used for real-time quantitative PCR.

| **Gene** | | **Primer Sequence** |
| --- | --- | --- |
| *GAPDH* | fwd | 5'-ATGGTGAAGGTCGGTGTGAA-3' |
|  | rev | 5'-AGTGGAGTCATACTGGAACA-3' |
| *DMT1* | fwd | 5'-AGGTGACACTATAGAATAGCCAGCCAGTAAGTTCAAGG-3' |
|  | rev | 5'-GTACGACTCACTATAGGGAGCTGTCCAGGAAGACCTGAG-3' |
| *FHC* | fwd | 5'-TGGAACTGCACAAACTGGCTACT-3' |
|  | rev | 5'-ATGGATTTCACCTGTTCACTCAGATAA-3' |
| *FLC* | fwd | 5'-CGTGGATCTGTGTCTTGCTTCA-3' |
|  | rev | 5'-GCGAAGAGACGGTGCAGACT-3' |
| *FPN1* | fwd | 5'-GGGTGGATAAGAATGCCAGACTT-3' |
|  | rev | 5'-GTCAGGAGCTCATTCTTGTGTAGGA-3' |
| *HAMP* | fwd | 5'-GGCAGACATTGCGATACCAAT-3' |
|  | rev | 5'-TGCAACAGATACCACACTGGGAA-3' |
| *IRP1* | fwd | 5'-GCGGATCCTACTGGGCCATCTTTCGGAT-3' |
|  | rev | 5'-GCTCTAGAGGCAGAGAGCTATGAGCGCATTCAC-3' |
| *IRP2* | fwd | 5'-AGGTGACACTATAGAATATGAAGAAACGGACCTGCTCT-3' |
|  | rev | 5'-GTACGACTCACTATAGGGAGCTCACATCCAACCACCTCT-3' |
| *TFR1* | fwd | 5'-CGCTTTGGGTGCTGGTG-3' |
|  | rev | 5'-GGGCAAGTTTCAACAGAAGACC-3' |
